## Supplementary Information for "Harvesting and amplifying gene cassettes confers cross-resistance to critically important antibiotics"

**Table S1**

| **ISOLATE** | **CTX-S/R** | **Transposon type** | **AMK-S/R** | **ST** | **3GC-R gene** | **AMC-R gene** | **PMQR gene** |
| --- | --- | --- | --- | --- | --- | --- | --- |
| BS51 | CTX-R | WT Transposon | R | 131 | blaCTX-M-15 | blaOXA-1 | aac(6')Ib-cr |
| BS52 | CTX-R | Promoter, RBS, ATG deletion | S | 131 | blaCTX-M-15 |  | aac(6')Ib-cr |
| BS53 | CTX-R | WT Transposon | R | 131 | blaCTX-M-15 | blaOXA-1 | aac(6')Ib-cr |
| BS54 | CTX-R |  |  | 38 | blaCTX-M-27 | blaTEM-1 |  |
| BS55 | CTX-R |  |  | 131 | blaCTX-M-27 |  |  |
| BS56 | CTX-R | C-T in promoter -10 | R | 131 | blaCTX-M-15 | blaOXA-1 | aac(6')Ib-cr |
| BS57 | CTX-R | WT Transposon | R | 131 | blaCTX-M-15 | blaOXA-1 | aac(6')Ib-cr |
| BS58 | CTX-R |  |  | 131 | blaCTX-M-27 |  |  |
| BS17 | CTX-R | WT Transposon | R | 131 | blaCTX-M-15 | blaOXA-1 | aac(6')Ib-cr |
| BS15 | CTX-R |  |  | 501 | blaCTX-M-15 |  |  |
| BS32 | CTX-R | WT Transposon | R | 38 | blaCTX-M-15 | blaOXA-1 | aac(6')Ib-cr |
| BS33 | CTX-R |  |  | 131 | ampC-32 | ampC-32 |  |
| BS19 | CTX-R | WT Transposon | R | 131 | blaCTX-M-15 | blaOXA-1 | aac(6')Ib-cr |
| BS34 | CTX-R | WT Transposon | R | 131 | blaCTX-M-15 | blaOXA-1 | aac(6')Ib-cr |
| BS36 | CTX-R |  |  | 131 | blaCTX-M-15 | blaTEM-1 |  |
| BS31 | CTX-R | WT Transposon | R | 131 | blaCTX-M-15 | blaOXA-1 | aac(6')Ib-cr |
| BS26 | CTX-R |  |  | 88 | ampC-42 | ampC-42 |  |
| BS1 | CTX-R | WT Transposon | R | 131 | blaCTX-M-15 | blaOXA-1 | aac(6')Ib-cr |
| BS9 | CTX-R |  |  | 73 | blaCTX-M-15 |  |  |
| BS37 | CTX-R | WT Transposon | R | 1193 | blaCTX-M-15, blaCMY-60 | blaOXA-1, blaCMY-60 | aac(6')Ib-cr |
| BS84 | CTX-R |  |  | 501 | blaCTX-M-15 |  |  |
| BS59 | CTX-R |  |  | 23 | ampC-42 | ampC-42 |  |
| BS60 | CTX-R |  |  | 95 | blaDHA-1 | blaDHA-1 | qnrB4 |
| BS61 | CTX-R | WT Transposon | R | 131 | blaCTX-M-15 | blaOXA-1 | aac(6')Ib-cr |
| BS62 | CTX-R | WT Transposon | R | 131 | blaCTX-M-15 | blaOXA-1 | aac(6')Ib-cr |
| BS63 | CTX-R |  |  | 1431 | ampC-42 | ampC-42 |  |
| BS83 | CTX-R |  |  | 501 | blaCTX-M-15 |  |  |
| BS65 | CTX-R |  |  | 131 | blaCTX-M-15 | blaTEM-1 |  |
| BS66 | CTX-R | WT Transposon | R | 131 | blaCTX-M-15 | blaOXA-1 | aac(6')Ib-cr |
| BS24 | CTX-R |  |  | 131 | blaCTX-M-15 | blaTEM-1 |  |
| BS67 | CTX-R |  |  | 131 | blaCTX-M-15 |  |  |
| BS79 | CTX-R | WT Transposon | R | 410 | blaCTX-M-15 | blaOXA-1 | aac(6')Ib-cr |
| BS68 | CTX-R |  |  | 73 | blaCTX-M-15 | blaTEM-1 |  |
| BS69 | CTX-R | WT Transposon | R | 12 | blaCTX-M-15 | blaOXA-1 | aac(6')Ib-cr |
| BS71 | CTX-R |  |  | 450 | blaDHA-1 | blaDHA-1 | qnrB4 |
| BS73 | CTX-R | WT Transposon | R | 131 | blaCTX-M-15 | blaOXA-1 | aac(6')Ib-cr |
| BS74 | CTX-R | WT Transposon | R | 131 | blaCTX-M-15 | blaOXA-1 | aac(6')Ib-cr |
| BS80 | CTX-R |  |  | 200 | ampC-42 | ampC-42 |  |
| BS75 | CTX-R | WT Transposon | R | 131 | blaCTX-M-15 | blaOXA-1 | aac(6')Ib-cr |
| BS76 | CTX-R | WT Transposon | R | 131 | blaCTX-M-15 | blaOXA-1 | aac(6')Ib-cr |
| BS77 | CTX-R |  |  | 963 | blaCMY-2 | blaCMY-2 |  |
| BS25 | CTX-R |  |  | 998 | blaCTX-M-15 |  |  |
| BS39 | CTX-R | WT Transposon | R | 131 | blaCTX-M-15 | blaOXA-1 | aac(6')Ib-cr |
| BS8 | CTX-R | WT Transposon | S | 131 | blaCTX-M-15 | blaOXA-1 | aac(6')Ib-cr |
| BS3 | CTX-R |  |  | 131 | blaCTX-M-15 |  |  |
| BS22 | CTX-R | WT Transposon | S | 131 | blaCTX-M-15 | blaOXA-1 | aac(6')Ib-cr |
| BS41 | CTX-R | WT Transposon |  | 73 | ampC-32 | ampC-32 |  |
| BS27 | CTX-R | WT Transposon | R | 131 | blaCTX-M-15 | blaOXA-1, blaTEM-1 | aac(6')Ib-cr |
| BS45 | CTX-R | WT Transposon | S | 131 | blaCTX-M-15 | blaOXA-1 | aac(6')Ib-cr |
| BS2 | CTX-R | WT Transposon | R | 131 | blaCTX-M-15 | blaOXA-1 | aac(6')Ib-cr |
| BS6 | CTX-R |  |  | 131 | blaCTX-M-27 |  |  |
| BS5 | CTX-R |  |  | 38 | blaCTX-M-14 | blaTEM-1 |  |
| BS4 | CTX-R |  |  | 131 | blaCTX-M-15 |  |  |
| BS20 | CTX-R | WT Transposon | S | 131 | blaCTX-M-15 | blaOXA-1 | aac(6')Ib-cr |
| BS14 | CTX-R | WT Transposon | S | 131 | blaCTX-M-15 | blaOXA-1, blaTEM-1 | aac(6')Ib-cr |
| BS11 | CTX-R |  |  | 450 | blaCTX-M-15 |  | qnrS1 |
| BS13 | CTX-R | WT Transposon | S | 405 | blaCTX-M-15 | blaOXA-1 | aac(6')Ib-cr |
| BS21 | CTX-R |  |  | 131 | blaCTX-M-27 |  |  |
| BS23 | CTX-R |  |  | 131 | blaCTX-M-27 |  |  |
| BS42 | CTX-R |  |  | 131 | blaCTX-M-15 |  |  |
| BS7 | CTX-R | WT Transposon | R | 131 | blaCTX-M-15 | blaOXA-1 | aac(6')Ib-cr |
| BS16 | CTX-R |  |  | 131 | blaCTX-M-27 |  | qepA |
| BS43 | CTX-R |  |  | 131 | blaCTX-M-15 |  |  |
| BS12 | CTX-R |  |  | 648 | blaCTX-M-15 |  |  |
| BS29 | CTX-R |  |  | 12 | blaCMY-2 | blaCMY-2 |  |
| BS18 | CTX-R |  |  | 2020 | blaCTX-M-15 | blaTEM-1 |  |
| BS28 | CTX-R | WT Transposon | R | 131 | blaCTX-M-15 | blaOXA-1 | aac(6')Ib-cr |
| BS44 | CTX-R | WT Transposon | R | 131 | blaCTX-M-15 | blaOXA-1 | aac(6')Ib-cr |
| BS10 | CTX-R |  |  | 12 | blaCMY-2 | blaCMY-2 |  |
| BS100 | CTX-S |  | R | 131 |  | blaTEM-1 |  |
| BS101 | CTX-R | WT Transposon | S | 1193 | blaCTX-M-15 | blaOXA-1 | aac(6')-Ib-cr |
| BS102 | CTX-S | WT Transposon | R | 38 |  | blaOXA-1 | aac(6')-Ib-cr |
| BS103 | CTX-R | WT Transposon | S | 8313 | blaCTX-M-15 | blaOXA-1 | aac(6')-Ib-cr |
| BS104 | CTX-R | C-T in promoter -10 | R | 131 | blaCTX-M-15 | blaOXA-1 | aac(6')-Ib-cr |
| BS105 | CTX-R | WT Transposon | R | 131 | blaCTX-M-15 | blaOXA-1 | aac(6')-Ib-cr |
| BS106 | CTX-R | WT Transposon | S | 131 | blaCTX-M-15 | blaOXA-1 | aac(6')-Ib-cr |
| BS107 | CTX-R |  | S | 58 | blaCTX-M-15 |  | qnrS1 |
| BS108 | CTX-R | WT Transposon | S | 131 | blaCTX-M-15 | blaOXA-1 | aac(6')-Ib-cr |
| BS109 | CTX-S | WT Transposon | S | 131 |  | blaOXA-1 | aac(6')-Ib-cr |
| BS110 | CTX-R |  | S | 131 | blaCTX-M-27 |  |  |
| BS111 | CTX-R |  | S | 131 | blaCTX-M-15 |  |  |
| BS112 | CTX-R | WT Transposon | R | 131 | blaCTX-M-15 | blaOXA-1 | aac(6')-Ib-cr |
| BS113 | CTX-R | WT Upstream OXA TRUNCATED | R | 131 | blaCTX-M-15 |  | aac(6')-Ib-cr |
| BS114 | CTX-R | WT Transposon | R | NOVEL | blaCTX-M-15 | blaOXA-1 | aac(6')-Ib-cr |
| BS115 | CTX-R | WT Transposon | R | 131 | blaCTX-M-15 | blaOXA-1 | aac(6')-Ib-cr |
| BS116 | CTX-S | WT Transposon | R | 131 |  | blaOXA-1 | aac(6')-Ib-cr |
| BS117 | CTX-R |  | S | 131 | blaCTX-M-14 | blaTEM-1 |  |
| BS118 | CTX-R | WT Transposon | R | 131 | blaCTX-M-15 | blaOXA-1 | aac(6')-Ib-cr |
| BS119 | CTX-S |  | R | 131 |  | blaTEM-1 |  |
| BS120 | CTX-R |  | S | 131 | blaCTX-M-15 |  |  |
| BS121 | CTX-S | WT Transposon | R | 131 |  | blaOXA-1 | aac(6')-Ib-cr |
| BS122 | CTX-R | WT Transposon | R | 131 | blaCTX-M-15 | blaOXA-1 | aac(6')-Ib-cr |
| BS123 | CTX-R | WT Transposon | R | NOVEL | blaCTX-M-15 | blaOXA-1 | aac(6')-Ib-cr |
| BS124 | CTX-R | WT Transposon | R | 131 | blaCTX-M-15 | blaOXA-1 | aac(6')-Ib-cr |
| BS125 | CTX-R | WT Transposon | R | 131 | blaCTX-M-15 | blaOXA-1 | aac(6')-Ib-cr |
| BS126 | CTX-R |  | S | 131 | blaCTX-M-27 |  |  |
| BS127 | CTX-R |  | S | 131 | blaCTX-M-27 |  |  |
| BS128 | CTX-S |  | R | 131 |  |  |  |
| BS129 | CTX-R | WT Transposon | R | 131 | blaCTX-M-15 | blaOXA-1 | aac(6')-Ib-cr |
| BS130 | CTX-R |  | S | 73 | blaCTX-M-15 |  |  |
| BS131 | CTX-R | WT Transposon | R | 131 | blaCTX-M-15 | blaOXA-1 | aac(6')-Ib-cr |
| BS132 | CTX-S |  | R | 410 |  | blaOXA-1 |  |
| BS133 | CTX-R | WT Transposon | S | 131 | blaCTX-M-15 | blaOXA-1 | aac(6')-Ib-cr |
| BS134 | CTX-R | WT Transposon | R | 131 | blaCTX-M-15 | blaOXA-1 | aac(6')-Ib-cr |
| BS135 | CTX-R |  | S | 69 | blaCTX-M-15 | blaTEM-1 | qnrS1 |
| BS136 | CTX-R |  | S | 131 | blaCTX-M-27 |  |  |
| BS137 | CTX-S | WT Upstream NO OXA | S | 131 |  |  | aac(6')-Ib-cr |
| BS138 | CTX-R | WT Transposon | R | 131 | blaCTX-M-15 | blaOXA-1 | aac(6')-Ib-cr |
| BS139 | CTX-R |  | S | 131 | blaCTX-M-15 |  |  |
| BS140 | CTX-R |  | S | 131 | blaCTX-M-15 | blaTEM-1 |  |
| BS141 | CTX-S | WT Upstream NO OXA | R | 131 |  |  | aac(6')-Ib-cr |
| BS142 | CTX-R |  | S | 131 | blaCTX-M-15 | blaTEM-1 | qnrB1 |
| BS143 | CTX-R | WT Transposon | R | 131 | blaCTX-M-15 | blaOXA-1 | aac(6')-Ib-cr |
| BS144 | CTX-R |  | S | 131 | blaCTX-M-1 | blaTEM-1 |  |
| BS145 | CTX-R | WT Upstream OXA and CR Separated | R | 131 | blaCTX-M-15 | (blaOXA-1) | aac(6')-Ib-cr |
| BS146 | CTX-R |  | S | 1486 | blaCTX-M-55 |  |  |
| BS147 | CTX-R |  | R | 1193 | blaSHV-12 | blaTEM-1 |  |
| BS148 | CTX-R | WT Transposon | R | 648 | blaCTX-M-15 | blaOXA-1, blaTEM-1 | aac(6')-Ib-cr |
| BS149 | CTX-R | WT Transposon | R | 131 | blaCTX-M-15 | blaOXA-1 | aac(6')-Ib-cr |
| BS150 | CTX-R | WT Transposon | R | 131 | blaCTX-M-15 | blaOXA-1 | aac(6')-Ib-cr |
| BS151 | CTX-R | WT Transposon | R | 131 | blaCTX-M-15 | blaOXA-1, blaTEM-1 | aac(6')-Ib-cr |
| BS152 | CTX-R |  | S | 88 | blaCTX-M-15 | blaTEM-1 | qnrS1 |
| BS153 | CTX-R |  | S | 131 | blaCTX-M-15 |  |  |
| BS154 | CTX-R |  | S | 12 | blaCTX-M-14 |  |  |
| BS155 | CTX-R |  | S | 1193 | blaCTX-M-55 | blaTEM-1 |  |
| BS156 | CTX-R |  | S | 10 | blaCTX-M-15 | blaTEM-1 |  |
| BS157 | CTX-S | WT Upstream OXA and CR Separated | R | 131 |  | blaOXA-1 | aac(6')-Ib-cr |
| BS158 | CTX-R | WT Transposon | R | 410 | blaCTX-M-15 | blaOXA-1 | aac(6')-Ib-cr |
| BS159 | CTX-R | WT Transposon | S | 131 | blaCTX-M-15 | blaOXA-1 | aac(6')-Ib-cr |
| BS160 | CTX-R |  | S | 131 | blaCTX-M-27 |  |  |
| BS161 | CTX-R |  | S | 69 | blaCTX-M-15 | blaTEM-1 |  |
| BS162 | CTX-R |  | S | 95 | blaCMY-2 | blaCMY-2 |  |
| BS163 | CTX-R | WT Upstream OXA TRUNCATED | S | 131 | blaCTX-M-15 |  | aac(6')-Ib-cr |
| BS164 | CTX-R | WT Transposon | R | 131 | blaCTX-M-15 | blaOXA-1 | aac(6')-Ib-cr |
| BS165 | CTX-R | WT Transposon | R | 131 | blaCTX-M-15 | blaOXA-1 | aac(6')-Ib-cr |
| BS166 | CTX-R |  | S | 131 | blaCTX-M-15 | blaTEM-1 |  |
| BS167 | CTX-R |  | S | 131 | blaCTX-M-15 |  |  |
| BS168 | CTX-R |  | S | NOVEL | blaCTX-M-15 | blaTEM-1 | qnrS1 |
| BS169 | CTX-R |  | S | 131 | blaCTX-M-15 | blaTEM-1 |  |
| BS170 | CTX-R | WT Transposon | R | 131 | blaCTX-M-15 | blaOXA-1 | aac(6')-Ib-cr |
| BS171 | CTX-R | WT Upstream OXA TRUNCATED | R | 131 | blaCTX-M-15 | blaTEM-1 | aac(6')-Ib-cr |
| BS172 | CTX-R |  | S | 131 | blaCTX-M-15 |  |  |
| BS173 | CTX-R | WT Transposon | R | 131 | blaCTX-M-15 | blaOXA-1 | aac(6')-Ib-cr |
| BS174 | CTX-R |  | S | 131 | blaCTX-M-15 |  |  |
| BS175 | CTX-R |  | S | 131 | blaCTX-M-15 |  |  |
| BS176 | CTX-R |  | S | 131 | blaCTX-M-15 |  |  |
| BS177 | CTX-S |  | R | 744 |  | blaTEM-1 |  |
| BS178 | CTX-S | WT Upstream OXA TRUNCATED | R | 648 |  |  | aac(6')-Ib-cr |
| BS179 | CTX-R |  | S | 131 | blaCTX-M-15 |  |  |
| BS180 | CTX-R |  | S | 131 | blaCTX-M-15 |  |  |
| BS181 | CTX-R |  | S | 141 | blaCTX-M-15 |  |  |
| BS182 | CTX-R |  | S | 5150 | blaCMY-2 | blaTEM-1, blaCMY-2 |  |
| BS183 | CTX-R |  | S | 131 | blaCTX-M-27 |  |  |

CTX, Cefotaxime; AMK, Amikacin; ST, Sequence Type; 3GC-R, 3GC-resistance; AMC-R, amoxicillin/clavulanate resistance; PMQR, Plasmid Mediated Quinolone Resistance.
